# No evidence of sentinel behaviour in a highly social bird based on an artificial set-up

**DOI:** 10.64898/2026.03.17.712373

**Authors:** Marta Marmelo, Liliana Silva, André Ferreira, Claire Doutrelant, Rita Covas

## Abstract

Sentinel behaviour occurs when individuals use raised positions to scan for predators while the rest of the group forages. Here, we investigated whether a colonial cooperatively breeding species that forages in large groups, the sociable weaver, *Philetairus socius*, displays sentinel behaviour. This behaviour has been reported in species with similar ecology, behaviour and foraging habits, (e.g. ground foraging in open habitats where aerial predators are common) and, hence, we expected that it could occur in sociable weavers. On the other hand, sentinel behaviour appears to be less common in species that live in very large groups. We used an experimental set-up consisting of an artificial feeding station and perches to assess occurrence of sentinel related behaviours: (i) perching events > 30s on an elevated position, (ii) head-movements and (iii) alarm calling. Birds were seldom observed perching while others fed, and those that did, perched for periods that were too short to be considered as sentinel behaviour (less than 5s on average). Our results suggest that this behaviour is uncommon or even absent in sociable weavers. We discuss whether other factors such as foraging in very large groups, or interspecific foraging associations might make sentinel behaviour less important in this species.

## Introduction

Individuals are considered to act as sentinels when they search for danger from an elevated vantage point, often rotating their heads for visual monitoring (Beauchamp, 2022). Moreover, to consider a behaviour as sentinel it needs to be coordinated, with individuals taking turns to conduct sentinel bouts (Bednekoff, 2015). To facilitate coordination, sentinels can produce a ‘watchman’s call’, to inform the other group members that they are on watch (Wickler, 1985), and also alarm calls when a predator is detected (Ridley, 2013).

Although studies have been conducted on species with variable group size, the coordination of sentinel behaviour has so far been mostly described in bird species living in stable and relatively small family groups and cooperative breeders (e.g. the Southern-pied babbler (Ridley, 2013)); see also (Table 1). It is currently poorly understood how common sentinel behaviour is in species living in larger groups.

**Table 1.**
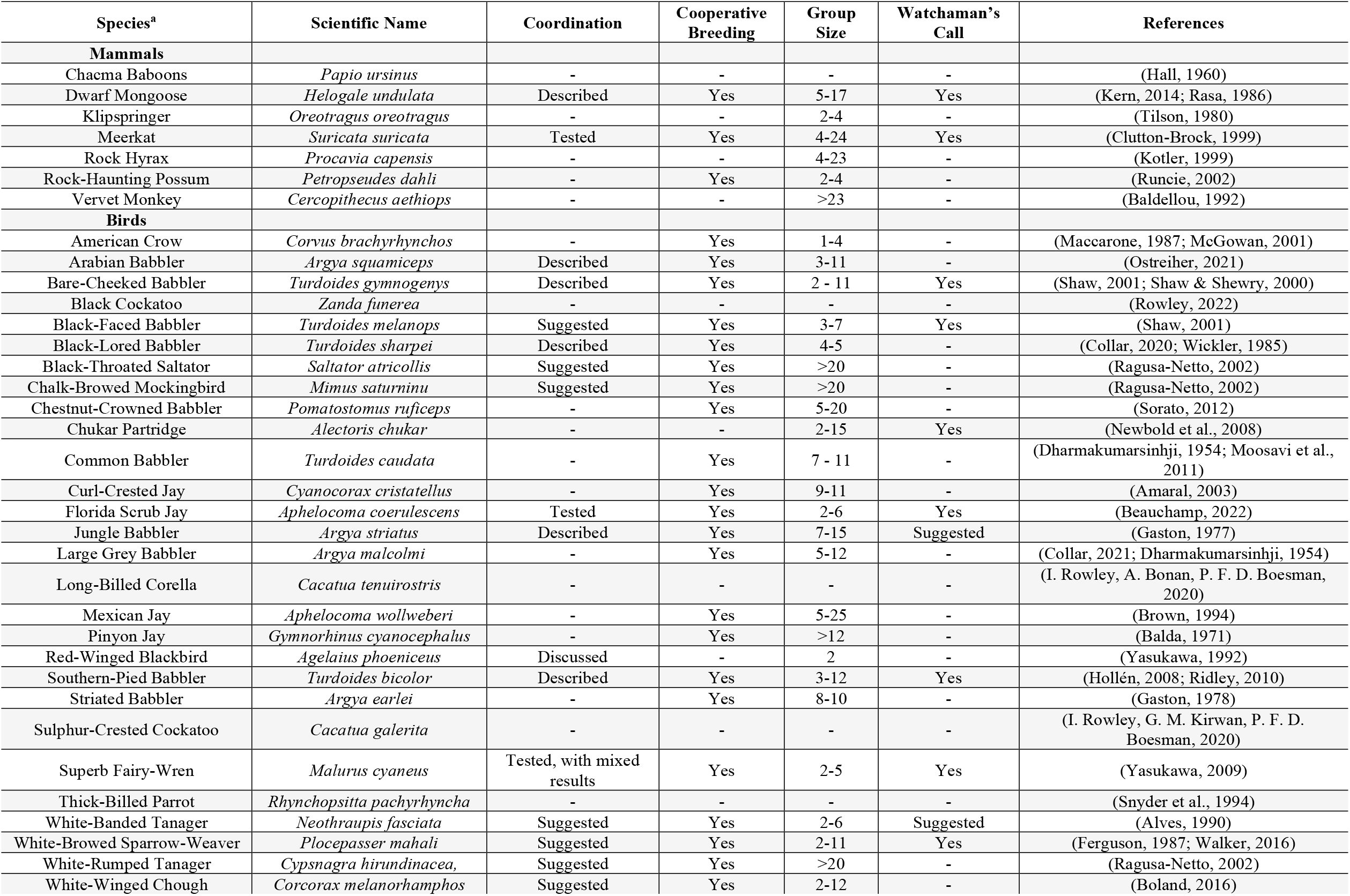
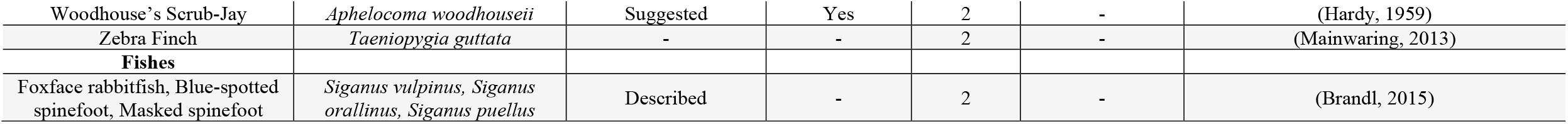
Species where sentinel like behaviour has been found, adapted from (Bednekoff, 2015) and supplemented with information on the occurrence of cooperative breeding, production of watchman calls, and group size when a sentinel is present. In most studies, observations of sentinel behaviour were conducted opportunistically. We use “-”when no information was found. **a**. Two bird species present in the original table were not included in our review: White-throated laughingthrush, (*Garrulax albogularis*) since we cannot access the book cited or any additional studies,and Northwestern crow (*Corvus caurinus*) since it is now considered a subspecies of the American crow (*Corvus brachyrhynchos*).

| Species<sup>a</sup> | Scientific Name | Coordination | Cooperative Breeding | Group Size | Watchaman's Call | References |
| --- | --- | --- | --- | --- | --- | --- |
| **Mammals** |  |  |  |  |  |  |
| Chacma Baboons | *Papio ursinus* | - | - | - | - | (Hall, 1960) |
| Dwarf Mongoose | *Helogale undulata* | Described | Yes | 5-17 | Yes | (Kern, 2014; Rasa, 1986) |
| Klipspringer | *Oreotragus oreotragus* | - | - | 2-4 | - | (Tilson, 1980) |
| Meerkat | *Suricata suricata* | Tested | Yes | 4-24 | Yes | (Clutton-Brock, 1999) |
| Rock Hyrax | *Procavia capensis* | - | - | 4-23 | - | (Kotler, 1999) |
| Rock-Haunting Possum | *Petropseudes dahli* | - | Yes | 2-4 | - | (Runcie, 2002) |
| Vervet Monkey | *Cercopithecus aethiops* | - | - | >23 | - | (Baldellou, 1992) |
| **Birds** |  |  |  |  |  |  |
| American Crow | *Corvus brachyrhynchos* | - | Yes | 1-4 | - | (Maccarone, 1987; McGowan, 2001) |
| Arabian Babbler | *Argya squamiceps* | Described | Yes | 3-11 | - | (Ostreiher, 2021) |
| Bare-Cheeked Babbler | *Turdoides gymnogenys* | Described | Yes | 2 - 11 | Yes | (Shaw, 2001; Shaw & Shewry, 2000) |
| Black Cockatoo | *Zanda funerea* | - | - | - | - | (Rowley, 2022) |
| Black-Faced Babbler | *Turdoides melanops* | Suggested | Yes | 3-7 | Yes | (Shaw, 2001) |
| Black-Lored Babbler | *Turdoides sharpei* | Described | Yes | 4-5 | - | (Collar, 2020; Wickler, 1985) |
| Black-Throated Saltator | *Saltator atricollis* | Suggested | Yes | >20 | - | (Ragusa-Netto, 2002) |
| Chalk-Browed Mockingbird | *Mimus saturninu* | Suggested | Yes | >20 | - | (Ragusa-Netto, 2002) |
| Chestnut-Crowned Babbler | *Pomatostomus ruficeps* | - | Yes | 5-20 | - | (Sorato, 2012) |
| Chukar Partridge | *Alectoris chukar* | - | - | 2-15 | Yes | (Newbold et al., 2008) |
| Common Babbler | *Turdoides caudata* | - | Yes | 7 - 11 | - | (Dharmakumarsinhji, 1954; Moosavi et al., 2011) |
| Curl-Crested Jay | *Cyanocorax cristatellus* | - | Yes | 9-11 | - | (Amaral, 2003) |
| Florida Scrub Jay | *Aphelocoma coerulescens* | Tested | Yes | 2-6 | Yes | (Beauchamp, 2022) |
| Jungle Babbler | *Argya striatus* | Described | Yes | 7-15 | Suggested | (Gaston, 1977) |
| Large Grey Babbler | *Argya malcolmi* | - | Yes | 5-12 | - | (Collar, 2021; Dharmakumarsinhji, 1954) |
| Long-Billed Corella | *Cacatua tenuirostris* | - | - | - | - | (I. Rowley, A. Bonan, P. F. D. Boesman, 2020) |
| Mexican Jay | *Aphelocoma wollweberi* | - | Yes | 5-25 | - | (Brown, 1994) |
| Pinyon Jay | *Gymnorhinus cyanocephalus* | - | Yes | >12 | - | (Balda, 1971) |
| Red-Winged Blackbird | *Agelaius phoeniceus* | Discussed | - | 2 | - | (Yasukawa, 1992) |
| Southern-Pied Babbler | *Turdoides bicolor* | Described | Yes | 3-12 | Yes | (Hollén, 2008; Ridley, 2010) |
| Striated Babbler | *Argya earlei* | - | Yes | 8-10 | - | (Gaston, 1978) |
| Sulphur-Crested Cockatoo | *Cacatua galerita* | - | - | - | - | (I. Rowley, G. M. Kirwan, P. F. D. Boesman, 2020) |
| Superb Fairy-Wren | *Malurus cyaneus* | Tested, with mixed results | Yes | 2-5 | Yes | (Yasukawa, 2009) |
| Thick-Billed Parrot | *Rhynchopsitta pachyrhyncha* | - | - | - | - | (Snyder et al., 1994) |
| White-Banded Tanager | *Neothraupis fasciata* | Suggested | Yes | 2-6 | Suggested | (Alves, 1990) |
| White-Browed Sparrow-Weaver | *Plocepasser mahali* | Suggested | Yes | 2-11 | Yes | (Ferguson, 1987; Walker, 2016) |
| White-Rumped Tanager | *Cypsnagra hirundinacea* | Suggested | Yes | >20 | - | (Ragusa-Netto, 2002) |
| White-Winged Chough | *Corcorax melanorhamphos* | Suggested | Yes | 2-12 | - | (Boland, 2016) |
| Woodhouse's Scrub-Jay | *Aphelocoma woodhouseii* | Suggested | Yes | 2 | - | (Hardy, 1959) |
| Zebra Finch | *Taeniopygia guttata* | - | - | 2 | - | (Mainwaring, 2013) |
| **Fishes** |  |  |  |  |  |  |
| Foxface rabbitfish, Blue-spotted spinefoot, Masked spinefoot | *Siganus vulpinus*, *Siganus orallinus*, *Siganus puellus* | Described | - | 2 | - | (Brandl, 2015) |

Here, we aimed to determine whether sociable weavers, *Philetairus socius*, a highly social colonial bird that lives and forages in large groups (ranging ca. 10-500 individuals), displays sentinel behaviour. In similarity to several sentinel species (Table 1), these weavers are cooperative breeders (Covas, 2008) and also cooperate in predator detection through alarm calls and in nest building (Maclean, 1973a; Marmelo, 2022). Sociable weavers predominantly forage in large flocks on the ground in open areas, with their heads down searching for seeds or invertebrates (Maclean, 1973c). Adults are mainly preyed on by an aerial predator, the Gabar goshawk (*Micronisus gabar*) (Maclean, 1973b), and while foraging sociable weavers regularly stop and raise their heads in vigilance (Marmelo, 2022). Hence, in this species, the presence of sentinels could allow group members to feed more efficiently. To assess if sociable weavers perform sentinel behaviour, we used an artificial set-up at feeding stations.

## Materials and Methods

### Study species and site

Sociable weavers are a small passerine endemic to Southern Africa (Maclean, 1973c). Artificial feeders, introduced since 2016 at five colonies (Ferreira, 2020) are set under a tree near each colony (ca. 120-200m away). These consist of four feeding boxes, each with four perches and four small standard plastic bird feeders (Fig. 1), being accessible three days/week.

**Fig. 1.**
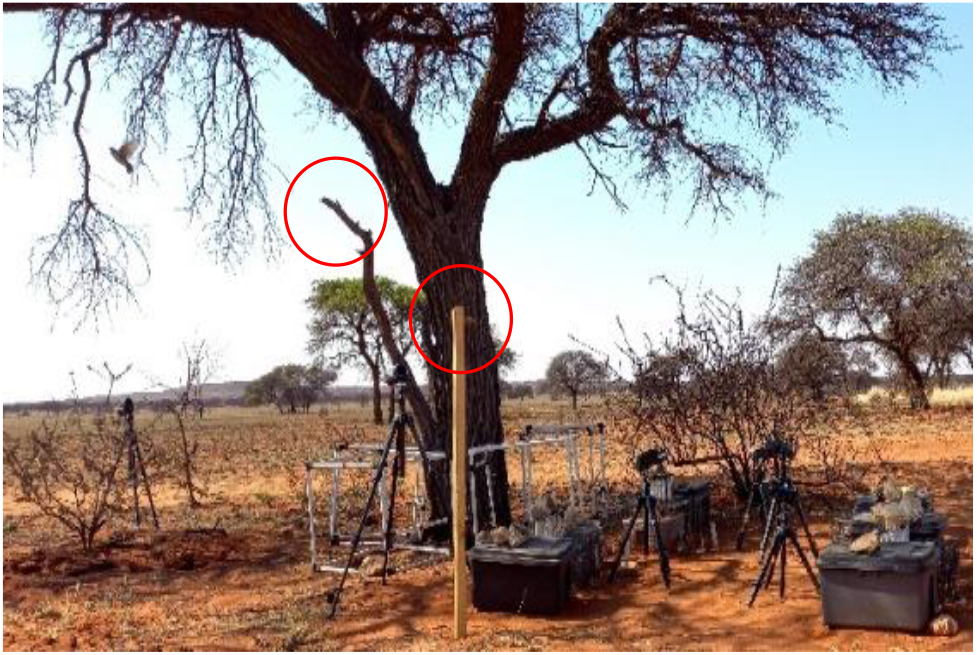
Example of a sentinel experimental set-up at a feeding station. Perches rounded in red (artificial on the centre - and natural on the left under tree).

### Data collection

The study was conducted in 2021, from September – November, at three colonies (ranging in size 13-56 individuals), where individuals were fully habituated to artificial feeders. We defined sentinel behaviour following previous studies (Table 1). We measured: (i) time spent on watch (i.e., perching on an elevated position such as a bush or small tree close to the foraging group), 30s being considered as a sentinel event, following Ridley (2010), (ii) occurrence of head movements (i.e., scanning) and (iii) alarm calls.

We placed one artificial and one natural perch (approximately 120 cm high) close to each other, and 90 cm away from the feeding station positioned next to a tree (Fig.1). Perches were positioned at the edge of the canopy tree to facilitate predator scanning. The artificial structure was made of cut wood. The natural one was a tree branch (provided at colonies 20 and 71; colony 11 had a natural branch that was known to be used as a perch by the birds (RC, personal observations). The birds were habituated to the structure’s presence for one to two months (depending on the colony) before we started collecting data.

We used video cameras to record the perches, between 7-12:00 am, corresponding to the main feeding time (Maclean, 1973c). At the same time, we also conducted direct observations from inside a tent (by MM), for two hours. The birds were habituated to human presence and their activity was not disturbed by the tent or cameras. From the videos we extracted perching and flying-away times, and which perches were used. All videos were manually analysed using BSPlayer™. In total, we obtained 9 days of video-recordings (three days per colony).

To complement this experiment, we conducted opportunistic observations in the field, under natural condition, when groups were foraging away from the colonies. Birds were observed with binoculars until they moved out of sight, either from a car (ca.15-50m away, on six occasions) or by following colonies on foot (on four occasions).

The STRANGEness of our test sample was evaluated as described by Webster and Rutz (2020). Our individuals are wild animals that were not raised by humans in specific conditions. Detailed breeding monitoring is conducted at our study site and some of the colonies have feeding stations nearby which are open every third day to collect data on social associations, but we have no reasons to think that these could affect the bird’s response to the treatments conducted in our experiment. These procedures have been approved by the Northern Cape Nature Conservation (permit FAUNA0059/2021) and the Ethics Committee of the University of Cape Town (permit 2020/2018/V22/RC/A1). Some birds could have been missing on the videos for random reasons, and by performing replicates of the experiment for three days at the artificial feeding stations, we think that we reduced potential sampling bias.

## Results & Discussion

We found no evidence of sentinel behaviour based on our artificial setup.

First, regarding the time perching on an elevated position as a surrogate of time spent on watch, we found that at two out three colonies, the birds never perched on the artificial structures provided. At one colony (colony 11), they used the existing tree branch, perching on 18 occasions. However, these occurrences were very rare and short, lasting on average four seconds (min=1s; max=12s) out of an average of 4 hours and 24 minutes recorded a day, during the 29 days of birds observed feeding. The duration of the perching events recorded in this study appears too short to be considered sentinel behaviour. It did not follow the minimum required of 30 seconds in this and in previous studies in species with similar ground foraging behaviour (Ridley, 2010; Walker, 2016). Regardless of the minimum required, it is important to note that sentinel behaviour usually lasts several minutes (Hollén, 2008; Kern, 2014).

Regarding the occurrence of head movements (i.e., scanning) and alarm calls, of the 18 short perching events mentioned above, 15 included head movements, and two included beak movement (associated with vocalisations). However clear head rotations (scanning), watchman’s calls and coordination in perching positions are described as supporting evidence of sentinel behaviour and were not found in sociable weavers.

Together these findings suggest that it is unlikely that sentinel behaviour is common in sociable weavers. However, our artificial set-up could have prevented us from detecting sentinel behaviour if sociable weavers were reluctant to perch on these artificial structures). This seems unlikely as these birds often perch on artificial structures (e.g., tripods and artificial feeders), were fully habituated to the feeding stations, and the perches provided were of a similar height to natural ones and set-up 1-2 months before the data collection. Furthermore, the observations conducted under natural conditions have not provided evidence of long perching times or other behaviours associated with sentinel.

The mean group size at the feeding stations during sentinel trials was small for this species (5±4.31 individuals, min=0; max=12, N=14 out of the 18 events), which means that the experiment might not have fully recreated normal foraging conditions and further systematic observations under natural conditions are needed to confirm these results. Nonetheless, smaller groups should in fact favour sentinel (Bednekoff, 2015).

Sentinels have been described in several group-living species but appear more common in species with smaller foraging groups (see Table 1). In our study area sociable weavers can forage in groups of >30 birds, and groups of several hundred birds are common in other areas. It is possible that in large foraging groups there are enough individuals being vigilant simultaneously to provide safety for the group (the ‘many eyes’ hypothesis, McNamara (1992)). In support of this mechanism, a previous study found that individual vigilance behaviour in sociable weavers, is observed at a rate of ca 34% of foraging time at the artificial feeding stations (Marmelo, 2022). In addition, we propose that an association with other species may reduce the need of vigilance behaviour. Fork-tailed drongos, *Dicrurus adsimilis*, commonly follow sociable weavers’ foraging flocks and produce watchman’s and alarm calls, leading to an increase in the weavers’ foraging time and decreased vigilance investment (Baigrie, 2014). We posit that the ‘many eyes’ of sociable weaver foraging flocks combined with their foraging associations with fork-tailed drongos might render sentinel behaviour less advantageous in this species than in others with similar foraging habits.

## Acknowledgements

The work would have not been possible without the contribution of several people working in the field, in particular field assistant Tshianeo Ndou and project manager Franck Theron. De Beers Consolidated Mines gave us permission to work at Benfontein Reserve. This study was supported by funding from the ERC to RC (EU, Consolidator grant 866489), ANR to CD (France, grant 19CE02-0014-01) and DST-NRF Centre of Excellence at the Fitzpatrick Institute of African Ornithology University of Cape Town (to RC). R.C. was funded by FCT (CEECIND/03451/2018).

